# Distinct functional roles for hopanoid composition in the chemical tolerance of *Zymomonas mobilis*

**DOI:** 10.1101/443473

**Authors:** Léa Brenac, Edward E.K. Baidoo, Jay D. Keasling, Itay Budin

## Abstract

Hopanoids are abundant membrane lipids found in diverse bacterial lineages, but their physiological roles are not well understood. The ethanol fermenter *Zymomonas mobilis* features the highest measured concentration of hopanoids, leading to the hypothesis that these lipids can protect against bacterial solvent toxicity. However, the lack of genetic tools for manipulating hopanoid composition *in vivo* has limited their further functional analysis. Because of polyploidy (> 50 genome copies per cell), we found that disruptions of essential hopanoid biosynthesis (*hpn*) genes in *Z. mobilis* act as genetic knockdowns, reliably modulating the abundance of different hopanoid species. Using a set of *hpn* transposon mutants, we demonstrate that both reduced hopanoid content and modified hopanoid head group composition mediate growth and survival in ethanol. In contrast, the amount of hopanoids, but not their polar group composition, contributes to fitness at low pH. Spectroscopic analysis of model membranes showed that hopanoids protect against several ethanol-driven phase transitions in membrane structure, including lipid interdigitation and bilayer dissolution. We propose that hopanoids act through a combination of hydrophobic and inter-lipid hydrogen bonding interactions to stabilize bacterial membranes against solvent stress.

**Graphical abstract:** 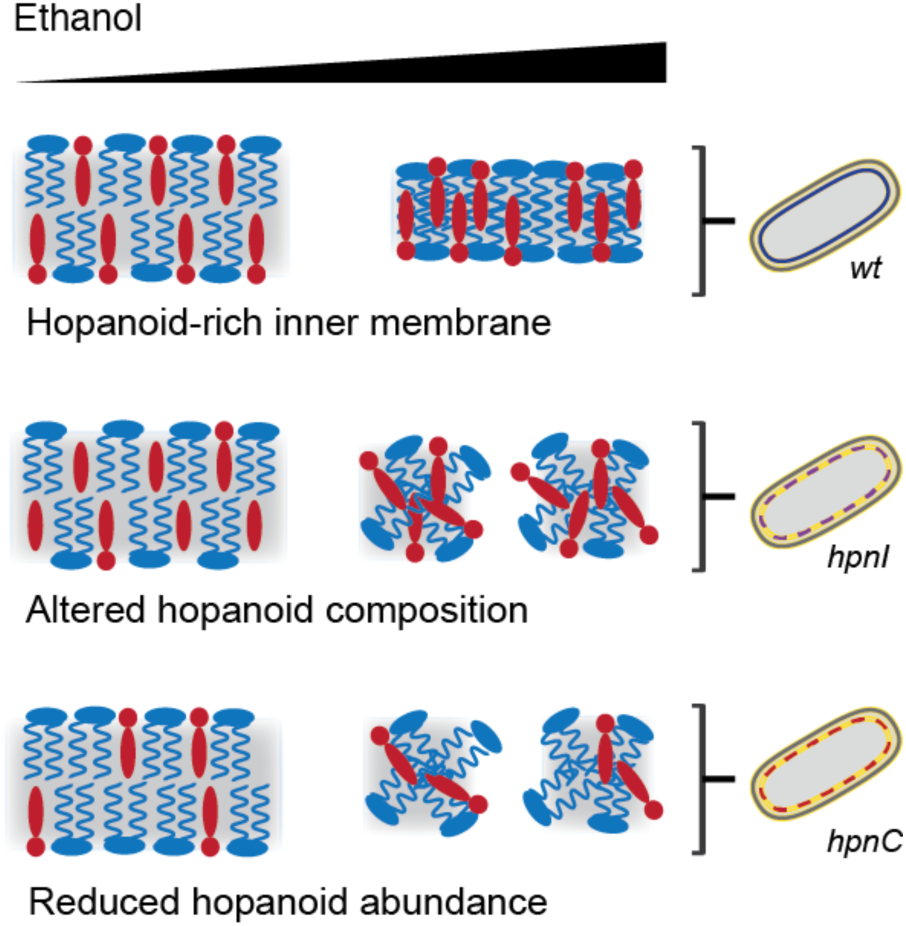

## Introduction

Triterpenoids are a class of cyclized lipids used by cells as bulk membrane building blocks. The canonical triterpernoids are eukaryotic sterols, in which squalene is oxidized before cyclization, leaving a hydroxyl to act as a polar head group. With very few exceptions (Pearson, Budin, & Brocks, 2003), bacteria instead synthesize hopanoids, in which squalene is first cyclized into hopene and then derivatized with polar groups (Ourisson & Rohmer, 1992). Hopanoids are thought to mimic the well-known effects of sterols on membrane ordering and permeability (Kannenberg, Blume, McElhaney, & Poralla, 1983; Poger & Mark, 2013; Saenz, Sezgin, Schwille, & Simons, 2012). Unlike sterols, however, they are generally present in complex mixtures of several derivatives, which could diversify their functions. They are also not as widespread, being absent in a majority of cultured bacterial strains (Rohmer, Bouvier-Nave, & Ourisson, 1984) and environmental samples (Pearson, Flood Page, Jorgenson, Fischer, & Higgins, 2007).

In bacteria with non-essential hopanoids, hopanoid biosynthesis (*hpn*) mutants have been used to identify functional roles related to chemical stress. In *Burkholderia cenocepacia*, hopanoid-free cells were found to be sensitive to low pH and detergents (Schmerk, Bernards, & Valvano, 2011), while in *Rhodopseudomonas palustris* they are more sensitive to both low and high pH (Welander et al., 2009) as well as bile salts (Welander et al., 2012). In *Bradyrhizobium diazoefficiens*, mutants lacking elongated hopanoids showed membrane defects and were sensitive to antibiotics, acid, and detergent (Kulkarni et al., 2015). The biophysical basis for these phenotypes has not been well studied, nor have physiological roles been characterized in bacteria where hopanoids are essential.

Because of its unusual glycolytic metabolism (Flamholz, Noor, Bar-Even, Liebermeister, & Milo, 2013), the alphaproteobacterium *Z. mobilis* produces ethanol approaching theoretical yields (Sprenger, 1996), leading to its adoption for industrial fermentation applications (Ming He et al., 2014). Native to palm wine fermenters (Swings & De Ley, 1977), *Z. mobilis* features a remarkable tolerance to ethanol, with reported growth in media containing up to 16% (v/v) ethanol (Lee, Skotnicki, Tribe, & Rogers, 1980). It also features a high abundance of hopanoids in its inner membrane – up to 50% of all lipids in the cell (Hermans, Neuss, & Sahm, 1991) – suggesting that these lipids could be important for its ability to withstand ethanol.

Like other solvents, ethanol alters the lipid packing (Vierl, Löbbecke, Nagel, & Cevc, 1994) and phase properties (Gurtovenko & Anwar, 2009) of model membranes. In principle, the condensing effect of hopanoids, analogous to eukaryotic sterols (Perzl et al., 1998; Saenz et al., 2012), could buffer against changes to membrane structure induced by solvents. However, in contrast to longer chain alcohols that fluidize membranes, the physicochemical effects of ethanol are complex, and include the induction of membrane phase transitions that increase membrane ordering (Vierl et al., 1994). Notably, in other ethanol-tolerant microbial fermentors (budding yeast), unsaturated fatty acids have been shown to mediate ethanol tolerance (Thomas, Hossack, & Rose, 1978; You, Rosenfield, & Knipple, 2003) and these lipids are thought to have opposing biophysical effects as hopanoids and other triterpenoids. In *Z. mobilis*, early lipid analyses using thin layer chromatography reported an increase in hopanoids during ethanol exposure (Bringer, Thomas, Poralla, & Sahm, 1985), but later work using modern mass spectrometry tools and transcriptomics showed no link between ethanol and hopanoid biosynthesis (Hermans et al., 1991; Moreau, Powell, Fett, & Whitaker, 1997) or *hpn* expression (Ming-xiong He et al., 2012), respectively. Therefore, it is still not established whether hopanoids contribute to the ethanol tolerance of *Z. mobilis*.

Testing physiological functions for hopanoid composition requires genetic tools to modulate their abundance and stoichiometry in cells. In *Z. mobilis*, a transposon-based gene disruption has previously been used to generate loss of function mutants to carry out systematic fitness analyses (Skerker et al., 2013). Surprisingly, this effort yielded transposon insertions in essential genes, e.g. those encoding for RNA/DNA polymerases. Such an effect was proposed to result from polyploid cells containing both mutant alleles, carrying antibiotic resistance markers, and functional ones, carrying essential genes. We hypothesized that this dynamic could reduce the copy number of biosynthetic genes, thereby functioning as genetic knockdowns and allowing for functional studies of hopanoid composition.

## Results

We first collected a redundant set of 25 Tn5 mutants of the *Z. mobilis* strain ZM4 covering annotated genes in the hopanoid biosynthesis pathway (Fig. 1A). Mutants with transposons at different positions in the same genes showed identical fitness profiles (Fig. S1), indicating that insertions acted as null alleles. When grown under antibiotic selection for the transposon, PCR analysis of transposon-inserted *hpn* genes showed the existence of both wild-type and mutated alleles (Fig. 1B), with varying apparent abundances. To confirm that heterozygosity was due to polyploidy, we measured chromosome numbers per cell directly using a qPCR-based approach (Fig. 1C). We found that *Z. mobilis* cells were highly polyploid, containing > 50 copies of two different genomic loci under our growth conditions. In contrast, a monoploid bacterium (*E. coli* in minimal medium) featured 1-2 genome copies per cell by this method. We quantified the transposon insertion rate for *hpn* mutants using high coverage genome sequencing, quantifying the number of reads matching to each allele (Fig 1D). The mutant allele frequency ranged from 31 to 100%, with non-duplicated genes early in the pathway featuring the lowest abundance and non-essential genes late in the pathway mimicking full knockouts (Table S1).

**Fig. 1:**
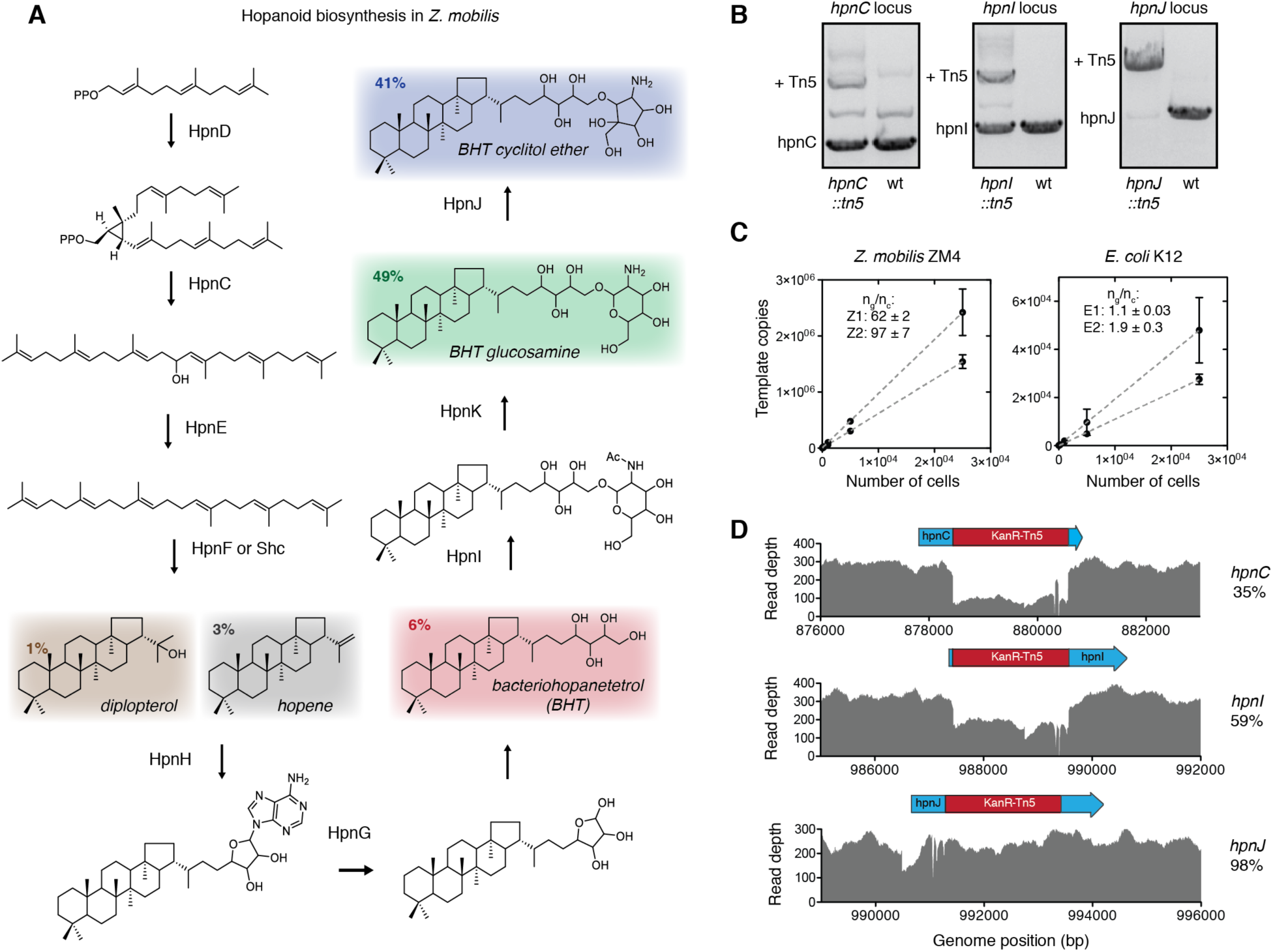
Genetic characterization of *hpn* transposon mutants in *Z. mobilis*. (**A**) Hopanoid biosynthesis in *Z. mobilis* occurs via a multistep pathway catalyzed by products of highly conserved *hpn* genes. Colored species indicate mature products and their reported (Hermans et al., 1991) abundance in wild type cells (as a % of total hopanoid weight content). (**B**) Characteristic colony PCRs showing the *hpn*::Tn5 strains contain both native *hpn* alleles and mutant *hpn*::Tn5 ones in varying amounts. In mutants early in the pathway (e.g. *hpnC*), the native allele is dominant over the mutant, but this is reversed for those late in the pathway (e.g. *hpnJ*). (**C**) Measurement of ploidy in *Z. mobilis* using qPCR of DNA extracted from counted cells, whose amplification was normalized with that of purified template standards. For *Z. mobilis* (left), this analysis yielded high genome copy numbers (60-100) for two different loci (Z1 and Z2). In contrast, *E. coli* grown in minimal media, where it is monoploid, showed a low copy number (1-2, E1 and E2) by the same method (right). Values represent mean +/-SEM, n=3. (**D**) Quantification of mutant allele abundance by deep sequencing of genomic DNA. Reads for each mutant were mapped to the genome sequence containing the transposon, and read coverage in the transposon was used to quantify the fraction of genome copies carrying the mutant allele. Shown are three examples that correspond to colony PCRs in panel (**B**); for each, a gene map with the transposon insertion is shown above the coverage plot. Quantification for the rest of the mutants is found in Table S1.

Since *hpn* mutants reduce the fraction of functional alleles in a cell, we hypothesized that they would reduce the abundance of the downstream product of each Hpn enzyme. To test this, we used a combination of mass spectrometry approaches to characterize the hopanoid composition of mutants grown under transposon selection. The products of squalene cyclization, hopene (Fig. 2A) and diplopterol (Fig. 2B), were measured by gas chromatography coupled to mass spectrometry (GC-MS) of silylated lipid extracts (Fig. S2A). Both showed decreased abundance in squalene synthase (*hpnC, hpnD*) mutants, consistent with their position upstream in the pathway. Lipids from *hpnF* and *shc* mutants, which exhibited almost complete disruption, showed distinct distributions of squalene cyclization products: the *shc* mutant accumulated less hopene, while the *hpnF* mutant accumulated less diplopterol. This difference suggests differing product preferences these SHCs. Lipids from the *hpnH* mutant exhibited higher hopene levels, consistent with the role of HpnH in converting hopene to adenosylhopane. Later genes in the pathway (*hpnG, hpnI, hpnK, hpnJ)* mildly altered levels of these products, possibly reflecting feedback regulation in the pathway, as have been well-characterized for sterol biosynthesis (Espenshade & Hughes, 2007).

**Fig. 2:**
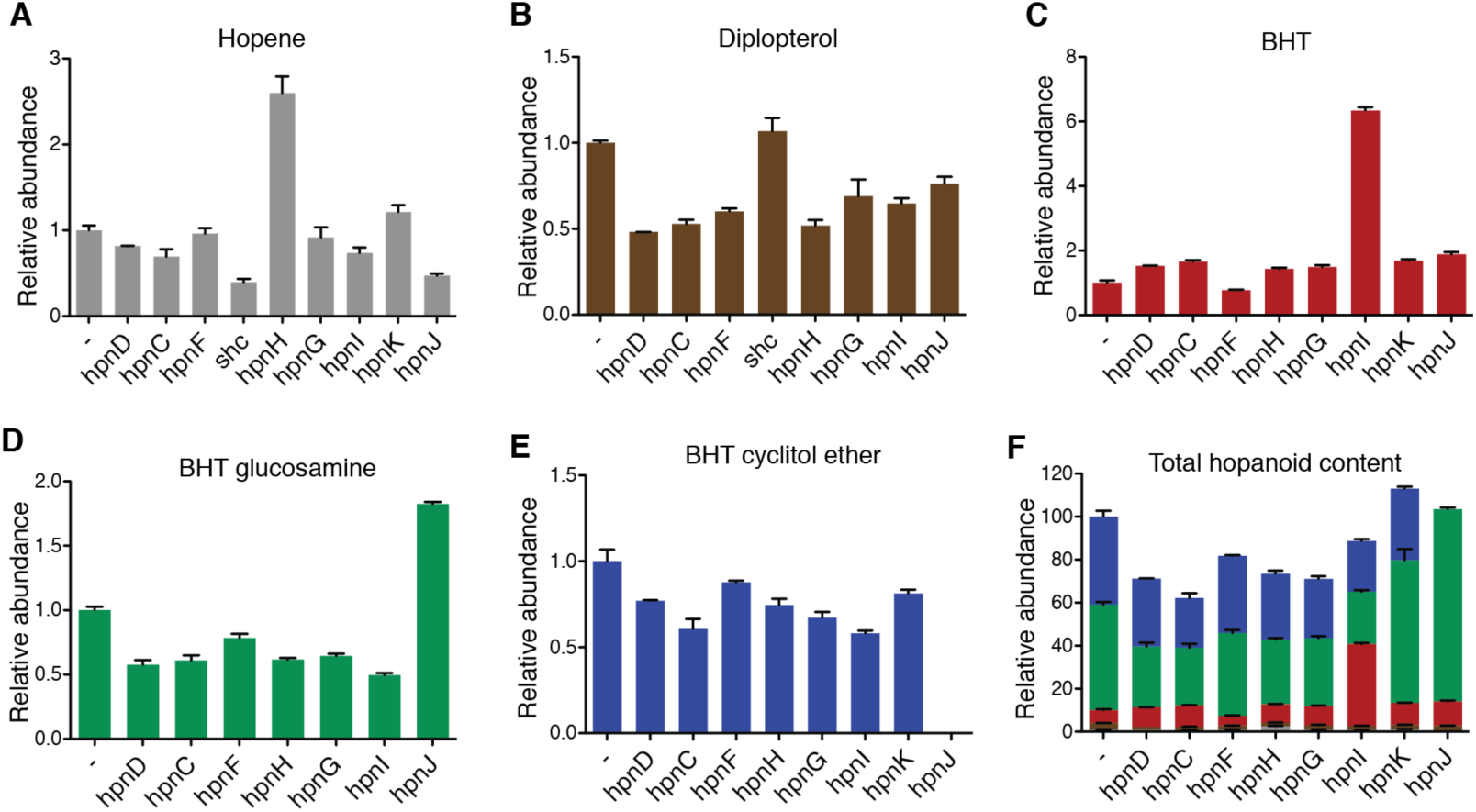
Composition of mature hopanoid species in *hpn* mutants or control cells (-). Cells were grown in RM media for 20-22 hours under kanamycin selection for the transposon and lipids were extracted as described in the Methods. Each species is color coded by their structures in Fig. 1a, and values are normalized to control cells. The abundance of hopene (**A**) and diplopterol (**B**) were measured by GC-MS of silylated lipid extracts, while BHT (**C**), BHT glucosamine (**D**) and BHT cyclitol ether (**E**) were measured by LC-APCI-TOF-MS of acetylated lipid extracts. Total hopanoids (**F**) are normalized by the absolute abundance of each species in wild type background as previously measured (Hermans et al., 1991). Error bars indicate mean +/-SD, n = 3 independent cultures.

We next analyzed the distribution of elongated hopanoids, which compose the majority of final hopanoid products in *Z. mobilis*, using liquid chromatography coupled to atmospheric pressure chemical ionization time of flight mass spectrometry (LC-APCI-TOF-MS) (Fig. 2C-E, Fig. S2B-C). Mutants targeting enzymes upstream of the elongated hopanoids all showed reduced abundance of all these products, but did not alter their stoichiometry. Instead, it was mutants of the three genes downstream of bacteriohopanetetrol (BHT) in the pathway (*hpnI, hpnK* and *hpnJ*) that altered elongated hopanoid distribution. The *hpnI* mutant exhibited an accumulation of BHT accompanied by a reduction of the BHT derivatives. Analyses of the *hpnK* mutant was hindered by the acetylation of BHT glucosamine during lipid derivatization. However, the increased ratio of BHT glucosamine to BHT cyclitol ether (2:1) we observed was consistent with its expected role as a deacetylase. In the *hpnJ* mutant, no BHT cyclitol ether could be detected, in agreement with it being near complete gene knock out (98% mutant alleles).

Lipid analysis confirmed the proposed biosynthetic route for hopanoids in *Z. mobilis* and showed that transposon disruptions act as functional knockdowns, resulting in changes in hopanoid abundance and distribution (Fig. 2F). We tested whether these changes modulate the physical properties of lipid bilayers using the anisotropy-based probe di-phenyl-hexatriene (DPH), a classic test of ordering in the hydrophobic membrane core (Suurkuusk, Lentz, Barenholz, Biltonen, & Thompson, 1976). Membranes prepared from extracted lipids with reduced hopanoid content showed significantly lower DPH anisotropy (Fig. 3), while those with altered head group composition showed a milder effect. This pattern is consistent with the hopene moiety – common to all hopanoids – being responsible for increasing hydrocarbon packing in the membrane, similar to the effect of sterols. All *Z. mobilis* compositions featured a higher anisotropy compared to membranes assembled from lipids extracted from *E. coli* MG1655, which lack hopanoids.

**Fig. 3:**
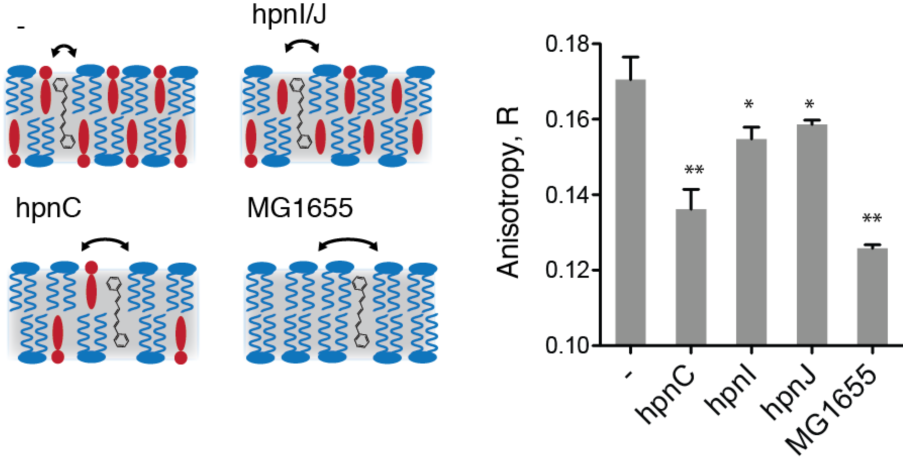
The dependence of bilayer order, as measured by DPH anisotropy, on hopanoid composition in liposomes. Liposomes were prepared using lipids extracted from early-stationary phase cultures for each *hpn* mutant, cells with intact hopanoids (-), or hopanoid-free *E. coli* (MG1655). Steady-state anisotropy of DPH incubated with the liposomes was measured as a unit-less ratio R = (I_A_ – I_B_) / (I_A_ + 2 I_B_), where I_A_ and I_B_ are the emission (430 nm) intensities parallel and perpendicular, respectively, to the polarization of the excitation (360 nm). Cartoons on the left diagram the compositional differences for each sample, as well as the relative motion of the probe within the bilayer’s hydrophobic core. In the diagrams, the blue and red species are phospholipids and hopanoids, respectively. Error bars indicate mean +/-SEM (n=3). Unpaired t-tests compare indicated lipid composition against the control (-): *, p < 0.05; **, p < 0.01.

We sought to use *hpn* mutants to assess the proposed physiological role for hopanoid composition in *Z. mobilis* ethanol tolerance. In standard rich media (RM), exponential growth rates of different mutants were correlated to their total hopanoid content, likely reflecting physiological roles for these lipids under non-stressed conditions (Fig. S3). We focused further investigation on a set of strains that sampled different aspects of hopanoid composition: reduced total hopanoid content (*hpnC* or *hpnD*), reduced accumulation of elongated hopanoids (*hpnI*), or absence of BHT cyclitol ether (hpnJ), comparing each with the behavior of a kanamycin-resistant control (Fig. 4A). For growth with 8% (v/v) ethanol, *hpn* mutants showed a significantly larger growth defect compared to the control (Fig. 4A-B). This effect was significant both for mutants with reduced hopanoid content and those with altered elongated hopanoids. A similar pattern was found for toxicity, measured by colony forming units, after acute (30 min) exposure to high ethanol concentration (20%), with each mutant featuring significantly reduced survival (Fig. 4C). Both hopanoid abundance and stoichiometry are thus functional for *Z. mobilis* ethanol fitness.

**Fig. 4:**
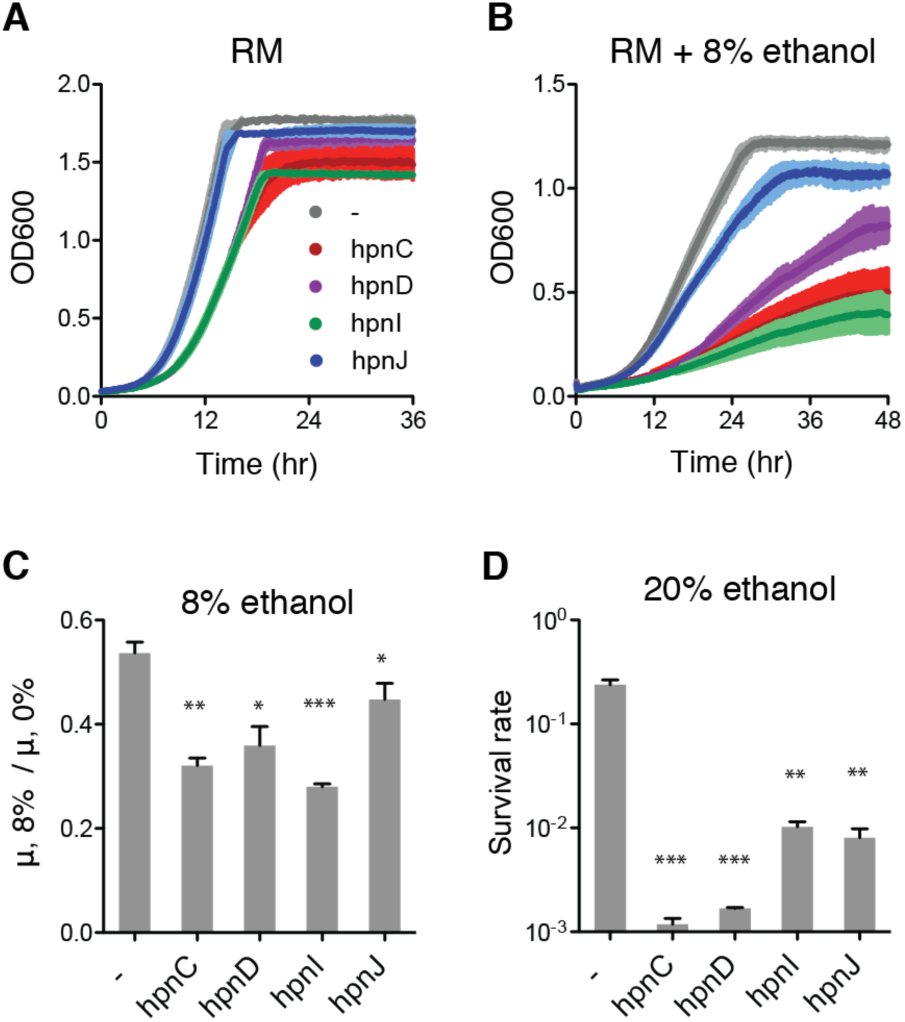
The effect of hopanoid knockdowns on *Z. mobilis* ethanol tolerance. A subset of *hpn* mutants sampling hopanoid composition were chosen for analysis. (**A**) Under standard conditions on rich medium (RM), *hpn* mutants grow similarly to control cells (-), which feature a kanamycin cassette in non-essential lysine exporter. In 8% v/v ethanol, however, *hpn* mutants show visible growth defects compared to control cells. (**B**) The relative growth rates in 8% vs. 0% ethanol (µ, 8% / µ, 0%) are significantly reduced for all *hpn* mutants compared to the control. (**C**) The *hpn* mutants also show significant increases in acute toxicity to ethanol, as measured by their survival rate after 30 min in 20% v/v ethanol. Error bars indicate mean +/-SEM, n = 3 independent cultures. Unpaired t-tests were used to assess significance against control cells (-): *, p < 0.05; **, p < 0.01; ***, p < 0.001.

In addition to ethanol, we considered other forms of chemical stress tolerance that would be characteristic of the native environment for *Z. mobilis*. Fermentation is characterized by acidic conditions, and *Z. mobilis* grows robustly between pH 3 and 4, matching the acidity of palm wine (Swings & De Ley, 1977). As discussed above, hopanoids have been known to increase ethanol tolerance in other bacteria in which they are not essential. Accordingly, we found that *hpn* mutants with reduced total hopanoid content (*hpnC* and *hpnD*) showed reduced growth rates at pH 3.9 (Fig. 5A) and survival at pH 2.5 (Fig. 5B).

**Fig. 5:**
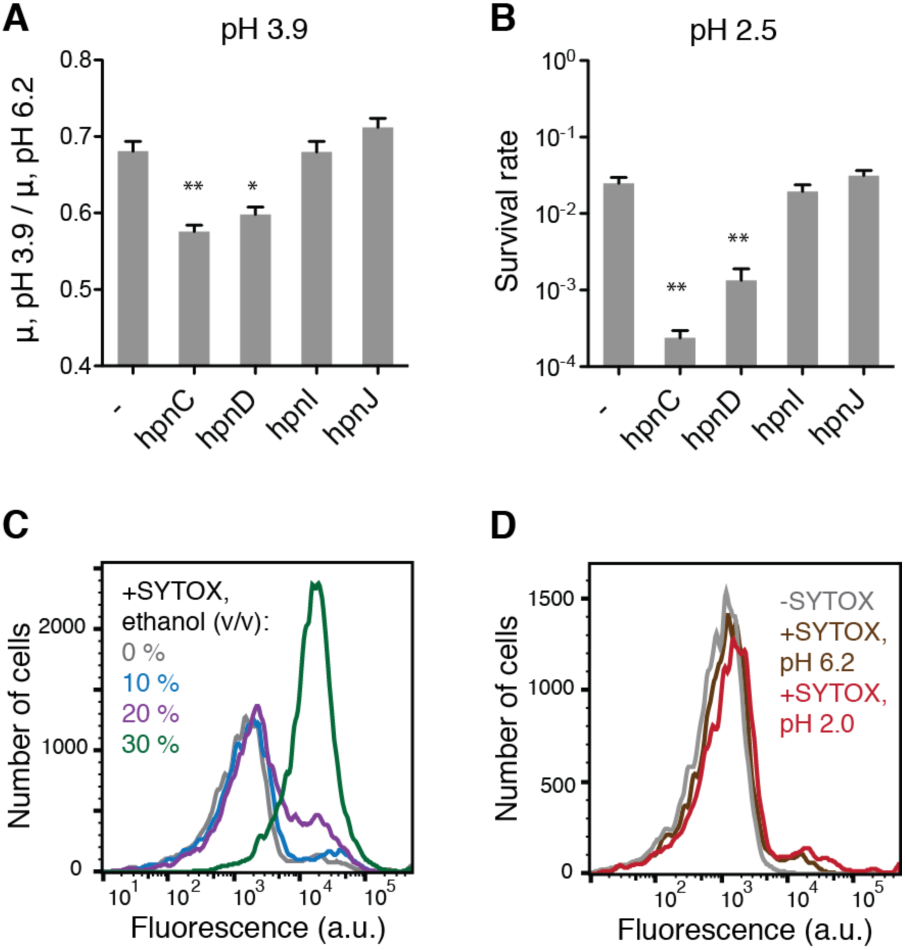
In contrast to ethanol, acid tolerance is dependent on hopanoid abundance, but not head group composition. (**A**) Mutants (*hpnC* and *hpnD*) with reduced hopanoid abundance show impairment in growth at pH 3.9 vs. pH 6.2, while those that affected hopanoid composition (*hpnI* and *hpnJ*) do not. (**B**) A similar pattern was observed for acute toxicity at high acidity (30 min at pH 2.5), as measured by colony forming units. Error bars indicate mean +/-SEM, n = 3 independent cultures. Unpaired t-tests were used to assess significance against control cells (-): *, p < 0.05; **, p < 0.01; ***, p < 0.001. (**C**) SYTOX staining of wild-type *Z. mobilis* cells with increasing ethanol concentrations shows that cell death between 20 and 30% (v/v) ethanol corresponds to a loss of membrane integrity. When permeabilization occurs, the polar dye can bind to cellular nucleic acids, leading to increased fluorescence. (**D**) In contrast, staining of cells in extreme acidity (pH 2.0) did not exhibit a loss in membrane integrity. Cells under these conditions showed no survival in physiological tests, suggesting that a loss of membrane structure is the cause of ethanol but not acid toxicity.

However, mutants that altered hopanoid composition, but not total abundance (*hpnI* and *hpnJ*), showed no significant change in acid tolerance. The presence of hopanoids is likely essential for *Z. mobilis* to thrive in acidic conditions, but the stoichiometry of head group species is less important as compared to that for ethanol toxicity.

The discrepancy the role of hopanoid stoichiometry in acid vs. ethanol could reflect different modes of lethality imposed by these chemical stresses. We hypothesized that while solvent exposure (e.g. ethanol) would lead to a loss of membrane structure, acid toxicity would occur by a different mechanism – such passive permeability of protons leading to cytoplasmic acidification – that is not dependent on loss of membrane integrity. To test this difference, we assayed membrane integrity using SYTOX staining. Cells incubated with ethanol exposure showed an increase membrane permeability at 20 and 30% v/v ethanol, corresponding to the toxicity threshold (Fig. 5C). However, those exposed to lethal acidic conditions (pH 2.0) did not show increased staining (Fig. 5D), indicating that acid does not lead to a loss in membrane integrity as ethanol exposure does.

To further explore the mechanisms by which hopanoids protect membranes against ethanol-mediated permeabilization, we carried out spectroscopic assays using liposomes assembled from *hpn* mutant lipid extracts. Because of its hydrophilicity, ethanol interacts with lipid membranes at the solvent-bilayer interface (Holte & Gawrisch, 1997). We therefore employed the fluorescent probe Laurdan, whose emission is sensitive to solvent access near the head group interface (Parasassi, Krasnowska, Bagatolli, & Gratton, 1998), to track changes to membrane structure in response to ethanol. As bilayers become disordered, they allow in more polar solvents, and Laurdan emission becomes red-shifted. This emission shift is quantified with a General Polarization (GP) ratio. Because Laurdan remains partitioned in lipid membranes independent of ethanol concentration (Zeng & Chong, 1995), changes in GP reflect changes in membrane ordering.

We found that Laurdan GP had a triphasic dependence on ethanol concentration, which track previously identified membrane structural changes (Fig. 6A-C). Initially, ethanol acts to disorder membranes, increasing head group spacing (bilayer expansion) and solvent accessibility, thereby reducing GP (regime I). At a critical concentration (∼10%), the trend is reversed as phospholipid acyl chains become interdigitated across the bilayer, a gel-like phase transition (Mou, Yang, Shao, & Huang, 1994; Simon & McIntosh, 1984) that increases GP (regime II). At a second critical concentration (>25%), Laurdan emission strongly red shifts, reflecting the dissolution of the bilayer structure at high ethanol concentrations that correspond to lethality in cells (regime III). The three regimes of the ethanol concentration vs. Laurdan GP relationship can thus be used to interrogate the specific structural effects on lipid membranes.

**Fig. 6:**
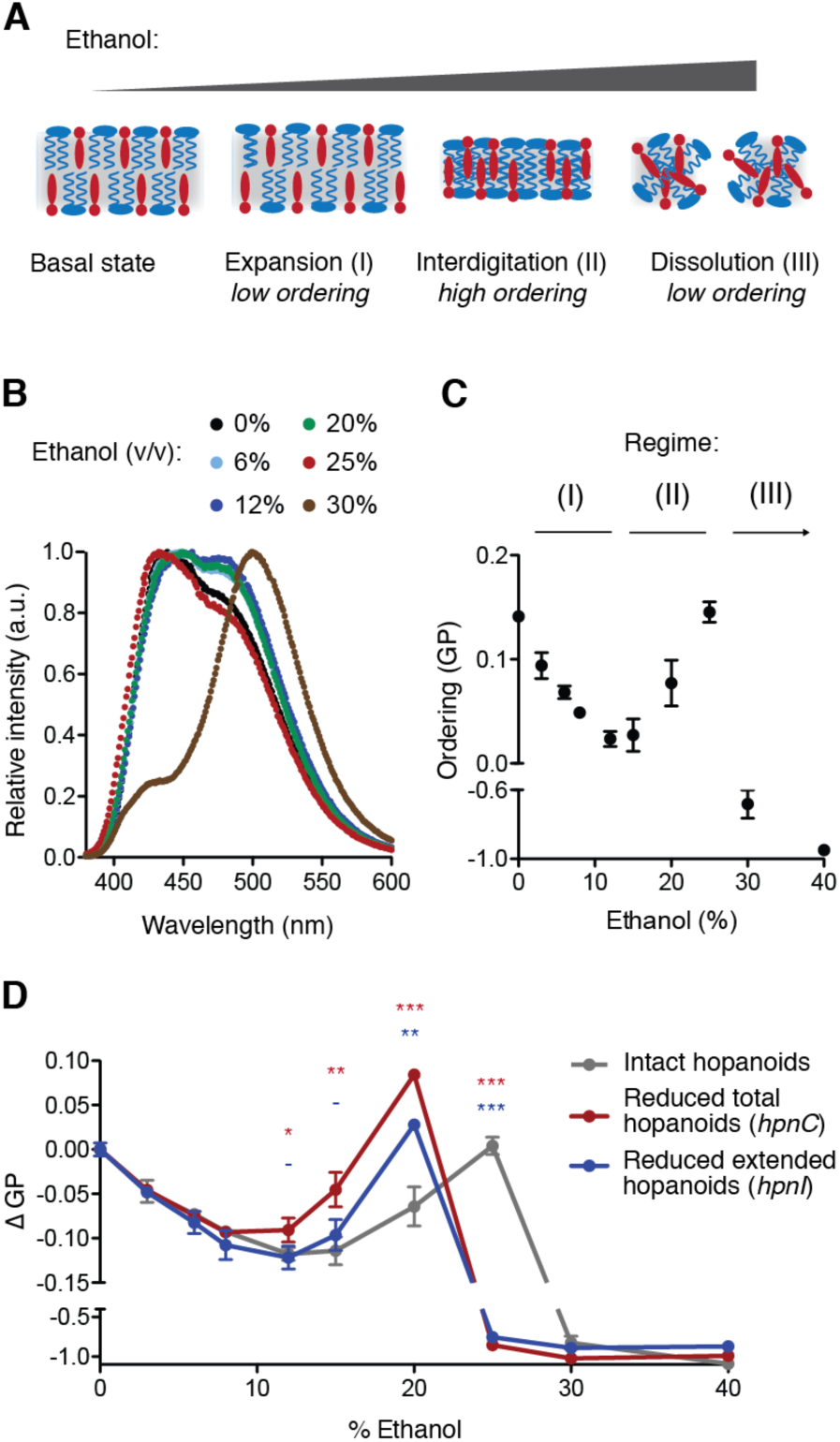
The hopanoid composition of *Z. mobilis*-derived liposomes mediates changes to bilayer structure induced by ethanol. (**A**) Cartoon showing the sequential phase transitions in membranes caused by increasing ethanol concentrations, including bilayer expansion (I), lipid interdigitation (II), and membrane dissolution (III). The blue and red species representing phospholipids and hopanoids, respectively. (**B**) Emission spectra of Laurdan in liposomes reconstituted from extracted *Z. mobilis* lipids with different amounts of ethanol. At low concentrations (0-10% v/v), ethanol red-shifts Laurdan emission, indicating more solvent access to the bilayer. At intermediate concentrations (10-25%), this effect is reversed due to an ordered membrane phase with interdigitated lipids. At high concentrations (> 30%), emission is strongly red-shifted, indicating solubilization of the membrane. (**C**) Quantification of membrane ordering, measured as Laurdan GP, and its triphasic dependence on ethanol concentration in membranes containing *Z. mobilis* lipids. Each of the three regimes (I, II, III) corresponds to a structural transition diagrammed in panel a. (**D**) The effect of mutant lipid compositions with hopanoid content (*hpnC*) or changes in its head group composition (*hpnI*) on changes to Laurdan GP with increasing ethanol concentrations. Mutant liposomes show a stronger increase in GP in moderate ethanol concentrations (12-20%), indicating a more pronounced interdigitated phase, as well as a reduced threshold for bilayer dissolution at high ethanol concentration. Error bars indicate mean +/-SD, n = 3 independent liposome preparations. Unpaired t-tests were used to assess significance against liposomes with intact hopanoids: *, p < 0.05; **, p < 0.01; ***, p < 0.001.

Consistent with our physiological data, hopanoid composition mediated the changes to membrane structure induced by ethanol (Fig. 6D). We compared the changes to Laurdan GP in liposomes made from extracts with mutants with reductions in total (*hpnC*) and extended (*hpnI*) hopanoids. Under basal conditions (0% ethanol), reductions in total (*hpnC*) and extended (*hpnI*) hopanoids led to decreased membrane ordering, with the latter having the larger effect (Fig. S4). This is in contrast with membrane ordering of the hydrophobic core measured with DPH (Fig. 3), likely reflecting the importance of the hopanoid polar head group in driving ordering at the solvent-bilayer interface, which Laurdan is sensitive to. This trend continued during bilayer expansion at 8% ethanol, which led to uniform decrease in Laurdan GP among all compositions. However, at higher ethanol concentrations (20%), ordering was significantly higher in hopanoid-depleted (*hpnC*) membranes, indicating a more pronounced interdigitated phase compared to membranes with wild type levels of hopanoids. At 25% ethanol, both *hpnC* and *hpnI* liposomes showed a dramatic drop in Laurdan GP, indicating a loss of membrane structure while control liposomes remained intact. Therefore, both hopanoid abundance and head group composition act to prevent membrane dissolution at high ethanol concentrations.

## Discussion

Although widespread among bacteria, hopanoids remain a poorly understood class of membrane lipids. To better understand their function, we sought a genetic system for modulating the composition hopanoids in *Z. mobilis*, a bacterium in which these lipids are both highly abundance and essential for growth. Because of its high polyploidy, we found that transposon mutants of essential *hpn* genes – when grown under selection – act as specific knockdowns of several steps in hopanoid biosynthesis. Using a set of *hpn* mutants, we characterized the effects of changes in hopanoid abundance and composition on the *Z. mobilis* physiology and the biophysical properties cell-derived membranes. Notably, we established that both the abundance of hopanoids and their polar head groups play key roles in dictating ethanol tolerance. A role for hopanoids in the ability of *Z. mobilis* to grow and survive in high ethanol concentrations has been hypothesized, but had been previously difficult to test. Because of the importance of this bacterium to industrial ethanol production, we expect that further engineering of hopanoid biosynthesis could be an avenue to tune its fermentation capabilities or to increase solvent tolerance in other, less-robust hosts.

We contrasted the role of hopanoids in ethanol tolerance with that of another chemical stress, acid. Unlike for ethanol, the ability for hopanoids to impart acid tolerance has been previously demonstrated in other bacteria. By being able to control both total abundance and the stoichiometry of different hopanoid species, we showed that only the former dictated survival at low pH. This finding suggests distinct mechanisms by which these two chemical stresses influence membranes, and therefore how this class of lipids can protect against them. Specifically, low pH does not disrupt the integrity of *Z. mobilis* membranes (Fig. 5D) while ethanol, as a solvent stress, does has such a structural effect.

Using a spectroscopic probe for membrane structure (Laurdan), we characterized the interplay of increasing ethanol concentrations and hopanoid composition on membrane structure. These experiments showed that *Z. mobilis*-derived membranes undergo ethanol-mediated structural changes previously identified in model bilayers. Hopanoids alter the magnitude of these effects and the ethanol concentrations at which they take place. Although the parameters for phase transitions in extracted membranes might not exactly correspond to those in intact cells, it is worth considering their physiological relevance. The toxicity threshold for ethanol in cells – around 20% v/v – corresponds to an interdigitated state we observed in liposomes. Although membranes are ordered in this state, they are also highly leaky (Zeng, Smith, & Chong, 1993), consistent with SYTOX staining of cells in 20% ethanol. Accordingly, the mutant with reduced hopanoid amount (*hpnC*) showed the strongest interdigitated effect, as measured by increase in Laurdan GP, and also the lowest survival under acute ethanol toxicity. Evaluating how hopanoids modulate ethanol-membrane interactions *in vitro* thus provides a mechanism by which they could act to increase physiological tolerance *in vivo*.

Hopanoids are commonly thought to mimic the effects of eukaryotic sterols, whose planar structures condense lipid bilayers, reducing the area per phospholipid, and acyl chain mobility (Yeagle, 1985). These effects on membrane ordering (Fig. 3) lower the permeability of polar molecules across the bilayer, including protons (Deamer & Nichols, 1989). A reduction in proton leakage into the cytoplasm provided by the presence of abundance hopanoids or sterols provides a straightforward mechanism for acid tolerance imparted by these lipids. During ethanol stress, hopanoids instead act to mediate the phase properties of membranes: bilayer expansion at low concentrations, interdigitation at medium concentrations, and dissolution at higher concentrations. All these transitions are likely resisted by the increased packing of lipids that triterpenoids impart. Notably, cholesterol has previously been shown to resist the interdigitated phase transition in lipid vesicles (Tierney, Block, & Longo, 2005). In this way the interplay of hopanoids and ethanol mimics that of sterols or other triterpenoids.

One way hopanoids differ from other triterpenoids is in their polar head groups, which feature elongated polyhydroxylated moieties. Elongated hopanoids make up ∼90% of all species in *Z. mobilis* and we found that their composition also mediates ethanol tolerance. We propose a very different functional analogue for these structures: di-acyl glycerol lipids modified by one or more sugar groups (glycolipids), which are highly abundant in plant thylakoid and in cell-wall less bacteria. Glycolipids promote membrane ordering through hydrogen bonding interactions between adjacent lipid head groups (Curatolo, 1987). The phase behavior of glycolipid-containing membranes can be mediated by changes in head group structure, such as the type of glucoside linkage in a disaccharide (Iwamoto, Sunamoto, Inoue, Endo, & Nojima, 1982). Sugar-modified lipids, including glycolipids, hopanoids, and ceramides, could act as a general mechanism to buffer membrane structure against chemical agents (solvents, detergents) or other physical stresses (temperature, shear forces) that alter hydrophobic interactions. Hopanoids, however, are unique in that they feature both hydrocarbon and polar structural elements that promote bilayer stability. It is likely that the combination of these features allows different hopanoids species to maintain membrane stability in response to the wide-range of environmental challenges experienced by bacterial cells.

## Experimental Procedures

### Cell growth and physiology

*Z. mobilis* strains were grown in rich medium (RM) (10 g/L yeast extract (Becton Dickinson), 2 g/L potassium phosphate dibasic, and 20 g/L glucose, pH 6.2) in culture tubes shaking at 200 rpm at 30°C, unless otherwise noted. For mutants, the medium was supplemented with kanamycin (100 µg/mL). For each culture, 5 mL of medium was inoculated from a glycerol stock and strain was allowed to grow for 2 to 3 days under selection. Cultures were then back-diluted into 10 mL fresh medium; the inoculation ratio was varied across ZM4 mutants (from 0.1% to 2.5% v/v) to account for growth rate variability and enable harvesting after 20-22 hours in early stationary phase. Cells were then harvested for lipid, DNA extraction, or toxicity tests. For lipid analysis, characteristic cultures of each mutant were also dried to completeness and massed, in order to correlate optical density to cell dry mass.

For *E. coli* experiments, K12 cells (MG1655) were grown in MOPS minimal medium pH 7.2 (Teknova) with 0.6% (w/v) succinic acid as the carbon source. Cells were grown to early stationary phase (OD ∼ 1.0) at 30°C in 10 mL cultures before being harvested for ploidy quantification and to stationary phase (OD ∼ 2.0) in 50 mL Luria Broth medium for lipid extractions.

For fitness analysis, *Z. mobilis* growth curves were recorded in 1 mL cultures in 24-well plates (Falcon) inoculated with 10 µL of overnight culture. The plates were sealed with a semi-permeable film (Thermo Fischer) and incubated at 30°C in a Synergy 4 plate reader (BioTek) with shaking and absorbance readings (600 nm) every five minutes, for 24 to 72 hours. For acute toxicity measurements, cells were spun down in early stationary phase and resuspended in RM (control) and RM supplemented with ethanol (20% v/v) or HCl (until pH 2.5). After incubation (30 min), cells were serially diluted and plated on RM agar plates in technical triplicates. Cell survival was measured by the ratio of colony forming units (CFUs) compared to the control condition. For assaying of permeabilization, cells were incubated with 30 nM of SYTOX Green Dead Cell Stain (Invitrogen) for 15 minutes. The fluorescence intensity of 100,000 cells was then measured on an Accuri flow cytometer (BD) using a GFP fluorescence filter set. Data was analyzed and plotted using FlowJo.

### Genetic analyses

The function of each *hpn* gene was deduced from their roles described by loss of function studies in other bacteria (Bradley, Pearson, Saenz, & Marx, 2010; Schmerk et al., 2015; Welander et al., 2012). For PCR and copy number measurements, 2 mL of cultures grown to early stationary phase and genomic DNA extracted with a Wizard Genomic DNA Purification Kit (Promega). Outside-in colony PCR of a subset of *hpn* genes was performed with PrimeSTAR DNA polymerase (TaKaRa) using the following primers:

~~~
*hpnC*: CCACATGATTTCCCGTTCAGCTTATG and CCGGTTCTTTCAGAAAAGCGGC
*hpnI*: GCCAAGGGCTTCGAGTTACTATAAACT and TGACTTTTTTAAGAGCCGCTACGGT
*hpnJ*: CGGAAGTGCGGTCACTTATCCC and ACTGAGAAGCCCTTAGAGCC
~~~

For quantifying transposon insertion rates, genomic DNA libraries were prepared using NEBnext DNA Library Prep Master Mix Set for Illumina (New England Biolabs) following the instructions for 400 bp insert libraries. Sequencing was performed using the 600-cycle Miseq Reagent Kit v3 (Illumina) on the MiSeq system (Illumina). For analysis, the online platform Galaxy (usegalaxy.org) was used, with alignments performed with the *Map with BWA-MEM* tool to ZM4 genomes to which a typical transposon sequence (KanR Tn5 with an inserted TagModule) had been juxtaposed. The resulting alignment file was opened with UGENE and the fragments aligned to the border of the transposon were examined to find the transposon insertion locus. The genome files were updated with actual transposon insertions and the alignment was repeated. The coverage depth was calculated with *plotCoverage* tool. To estimate the transposon insertion rate, the mean coverage depth over the whole genome was computed and compared to the mean coverage depth of a 500 bp conserved region inside the transposon’s KanR cassette.

Quantification of ploidy was performed using qPCR as previously described (Pecoraro, Zerulla, Lange, & Soppa, 2011). Briefly, cells were harvested at early stationary phase and serially diluted. A hemocytometer was used to count the density of cells in the dilutions, and solutions containing discrete number of cells were then prepared. From these, genomic DNA was quantitatively extracted using phenol:chloroform extraction after cell lysis, as previously described (Thompson et al., 2018). Separately, standards containing the two targeted loci were amplified from *Z. mobilis* (Z1, Z2) and *E. coli* genomic DNA (E1, E2) using the following primers:

~~~
Z1 standard: TTGTGTTAAAGGGAGCGAGAC and AGACATATGTCCACGGACTTCTAA
Z2 standard: AATTTCAAAGGGGCCTTAGCT and TTAAAGACCCTCCTACTGCTTGT
E1 standard: ATTGCCCATAAACGAGAATACC and GGAAGCTTGGATCAACCG
E2 standard: TAACGCGTCGTCTGAAACC and TAGACGCCAGCATGTTCGTAA
~~~

The single-product amplification of the resulting standards, 1 KB in length, was confirmed by agarose electrophoresis, which was then purified by a PCR cleanup kit (Qiagen). The concentration of the standard was quantified by absorbance at 260 nm and then serially diluted to prepare a standard curve. PCR reactions for both gDNA samples and the standards was prepared with SYBR Green Master Mix (Invitrogen), using primer pairs that amplified a ∼250-bp insert in the standards above:

~~~
Z1 analysis: TAAACAGCCTTTAAAATTTTGTGTTCTTG and
GTTTATTTTCGGTATTTTTGACGCAG
Z2 analysis: ATTTCGATATTGTTCTGCAGTATCG and
TACTCAAGACCTTCAATATTGTTTTAACAT
E1 analysis: AAACTGGATACCCGCCTG and CCTTTTGACGTCATCATCATTG
E2 analysis: ATCCGTGATGAAGTGGGG and AATGCAGTAACCACAGCC
~~~

Amplification was measured on a StepOnePlus Real-Time PCR System (Applied Biosystems). Quantification of gDNA abundance relative to the standard curve was performed using the ΔC_T_ method.

### Lipid analyses

The lipid extraction protocol used in this study was adapted from Bligh and Dyer (Bligh & Dyer, 1959). Cell pellets from 10 mL of culture were resuspended in a 400 microliters of water and sonicated for 10 minutes at 50°C. Chloroform and methanol (1:1, 5 mL) were added and the monophasic extraction mixture was bath sonicated for 1 hour at 50°C and then left to shake for 3 hours at 37°C. Water (2.5 mL) was then added and complete separation of the organic (chloroform) and water/methanol phases was achieved through centrifugation. The organic phase was collected and evaporated to dryness under N_2_ flux.

For analysis of hopene and diplopterol by GC-MS, dry lipids were silylated in 150 µL of N,O-Bis (trimethylsilyl) trifluoroacetamide (BSTFA kit, Supelco) and 150 µL of pyridine for 2 hours at 50°C. The solvent was then evaporated under N_2_ flux. The product was then extracted in hexane (100 µL) and filtered (Nylon 0.2 µm 500 µL centrifugal filters, VWR) before analysis. Gas chromatography was performed on an HP 6890 Series GC-System (Hewlett Packard) equipped with a DB-5MS capillary column (30 m length x 0.25 mm internal diameter, 25 µm film; J&W). 1 µL of sample was injected at a temperature of 250°C and the GC was run in splitless mode with a Helium (carrier gas) flow rate of 1 mL/min at a pressure of 9.36 psi. The temperature program of the oven was as follows: held at 80°C for 1 min after injection, heated to 280°C at 20°C/min, held at 280°C for 15 min, heated to 300°C at 20°C/min and held for 7 min. The column was coupled to a HP 5973 Mass Selective Detector (Hewlett Packard). The source and quad temperature were set to 300°C and 200°C, respectively. The solvent delay was set to 9 min. For quantification of C_30_ hopanoids, the SIM mode (191 *m/z* for hopene and 131 *m/z* for diplopterol) was used with a dwell time of 100 ms and 3 cycles/s. Data collection and processing were performed with HP Enhanced ChemStation software (Agilent Technologies).

For analysis of elongated hopanoids, lipid extraction was performed as described above, with a known amount (100 µg) of an internal standard (5α-pregnane-3β,20β-diol) added to the extraction mixture. The following derivatization procedure was adapted from Talbot et *al.* (Talbot, Summons, Jahnke, & Farrimond, 2003) The dried chloroform extract was acetylated in 3 mL of acetic anhydride/pyridine (1:1 v/v) at 50°C for 1 hour and left stand overnight at room temperature. The solvent was then evaporated under N_2_ flux and the acetylated lipids were dissolved in 250 µL of methanol/isopropanol (6:4 v/v). The samples were centrifuged for 10 minutes at maximum speed and transferred to glass vials. Liquid chromatography was performed with a Kinetex XB-C18 column (100 mm length x 2.1 mm internal diameter, 2.6 µm particle size; Phenomenex) at 50°C, with a 1200 Series HPLC system (Agilent Technologies). The injection volume was 3 µL and separation was achieved with a flow-rate of 0.45 mL/min and the following mobile phase gradient profile: 20% to 42% B in 2.2 min, held at 42% B for 3.6 min, 42% to 100% B in 0.1 min, held at 100% B for 2.1 min, 100% to 20% B in 0.1 min, held at 20% B for 4.3 min (where solvent A is composed of 30% isopropanol/40% methanol/30% water, and solvent B of 90% isopropanol/10% water; HPLC grade, Honeywell Burdick & Jackson). The HPLC system was coupled to an Agilent Technologies 6210 time-of-flight mass spectrometer (LC-TOF MS), with 1/5 post-column split, via a LAN card. Nitrogen was used as nebulizing (6 L/min) and drying (300°C, 30 lb/in^2^) gases. The vaporizer temperature was 250°C. The corona was set to 7 µA. Atmospheric pressure chemical ionization (APCI) was conducted in the positive ion mode with a capillary voltage of 3000 V. Full-scan mode (100–1500 *m/z*) at 1 spectrum/s was used for MS experiments. TOF-MS was calibrated with Agilent APCI TOF tuning mix. Data acquisition and integration were performed by the MassHunter Workstation (Agilent Technologies) and MassHunter Qualitative Analysis (Agilent Technologies), respectively. The targeted ions are presented in Table S2; quantified analytes were normalized to the internal standard in order to correct for variations in ionization strength.

### Biophysical analyses

Liposomes were prepared from lipids extracted from early stationary phase cells (100 mL) using a scaled-up version of the procedure described above. Upon complete drying of the organic phase under vacuum, the extracted lipids were massed and solubilized in chloroform at 1 mg/mL. For each liposome preparation, 300 µL of the lipid solution was transferred to a borosilicate test tube and dried with Argon. The resulting film was further dried for 1 hour under vacuum. Liposomes were formed by resuspending the film in buffer (20 mM HEPES, pH 7.0), followed by overnight incubation with gentle agitation. Liposome formation was confirmed by microscopy of the samples labeled with Rhodamine 6G, a membrane dye. Before analysis, liposomes were extruded to 100 nm with a miniextruder (Avanti Polar Lipids).

For DPH experiments, liposomes were incubated with 0.5% (v/v) of a 5 mM DPH (Sigma) stock solution in ethanol. Samples were then incubated for at least one hour under gentle agitation. Steady-state anisotropy was measured using a Fluorolog spectrofluorometer (Horiba) with automated polarizers; excitation was set to 360 nm, and emission at 430 nm. For Laurdan experiments, liposomes were incubated with 1.0% (v/v) of a 1 mg/ml stock solution of Laurdan (Invitrogen) in DMSO. Laurdan emission was measured from 380 to 600 nm upon excitation at 365 nm. Laurdan GP was calculated as the ratio of integrated emission intensities at two different ranges of wavelengths: GP = (I_420-440nm_ – I_490-510nm_) / (I_420-440nm_ + I_490-510nm_).

## Acknowledgments

Dr. Jeffrey Skerker kindly provided strains and technical assistance. Tristan de Rond, Patrick Shih, Mitchell Thompson, and Meytal Higgins provided helpful discussions. This work was part of the DOE Joint BioEnergy Institute (http://www.jbei.org) supported by the U. S. Department of Energy, Office of Science, Office of Biological and Environmental Research, through contract DE-AC02-05CH11231 between Lawrence Berkeley National Laboratory and the U. S. Department of Energy. This work was also supported by National Science Foundation grants MCB-1442724 and MCB-1715681 to J.D.K. The authors declare no conflicts of interest.

## Author contributions

L.B., J.K.D., and I.B. conceived the study and designed the experiments. L.B. and E.E.K.B. performed lipid analysis. L.B. and I.B. performed all other experiments. All authors discussed the results and wrote the paper.

## Abbreviated Summary

A set of knockdown, transposon mutants was used to analyze the physiological functions of hopanoid composition in the ethanologenic bacterium *Zymomonas mobilis*. Both the abundance of hopanoids and their stoichiometry mediated growth and survival in ethanol-rich medium. Liposome-based experiments showed that hopanoids mediate the concentrations at which ethanol-induced transitions in membrane structure occur, providing a mechanism for their protective effects.

## Supporting Information

**Table S1:** *Z. mobilis* ZM4 mutants used in this study. Insertion sites provide the genome location of the repeat region, the sequence immediately before the transposon insertion; the same sequence is repeated at the end of the transposon. Tn5 frequency is the fraction of alleles containing the transposon, as measured by deep sequencing.

| Gene code | Gene symbol | Gene product | Insertion site (bp) | Direction of Tn5 | Tn5 frequency (%) |
| --- | --- | --- | --- | --- | --- |
| *hpnC* | ZMO0869 | squalene synthase | 878429-878437 | forward | 31 |
| *hpnD* | ZMO0870 | squalene synthase | 878952-878960 | reverse | 35 |
| *hpnF* | ZMO0872 | squalene hopene cyclase | 882157-882165 | reverse | 99 |
| *shc* | ZMO1548 | squalene hopene cyclase | 1579153-1579161 | reverse | 100 |
| *hpnG* | ZMO0873 | phosphorylase | 883206-883214 | forward | 43 |
| *hpnH* | ZMO0874 | radical SAM protein | 884303-884311 | reverse | 25 |
| *hpnI* | ZMO0972 | glycosylase | 987434-987442 | forward | 59 |
| *hpnJ* | ZMO0973 | radical SAM protein | 989161-989169 | forward | 98 |
| *hpnK* | ZMO0974 | polysaccharide hydrolyase | 990783-990791 | reverse | 92 |
| **Control** | ZMO0047 | lysine exporter protein | 45590-45598 | forward | 100 |

**Table S2:** Base peaks of targeted analytes for LC-APCI-TOF-MS analysis of elongated hopanoids used in this study. Designated acetylated structures for the compounds are shown in Fig. S2c.

| Compound | Acetylated structure | Base peak | Corresponding ion |
| --- | --- | --- | --- |
| BHT | I | 655.50 *m/z* | [M+H-CH<sub>3</sub>COOH]<sup>+</sup> |
| BHT ether | II | 1002.62 *m/z* | [M+H]<sup>+</sup> |
| BHT glucosamine | III | 1002.62 *m/z* | [M+H]<sup>+</sup> |
| 5α-pregnane-3β,20β-diol (standard) | IV | 345.28 *m/z* | [M+H-CH<sub>3</sub>COOH]<sup>+</sup> |

**Fig. S1:**
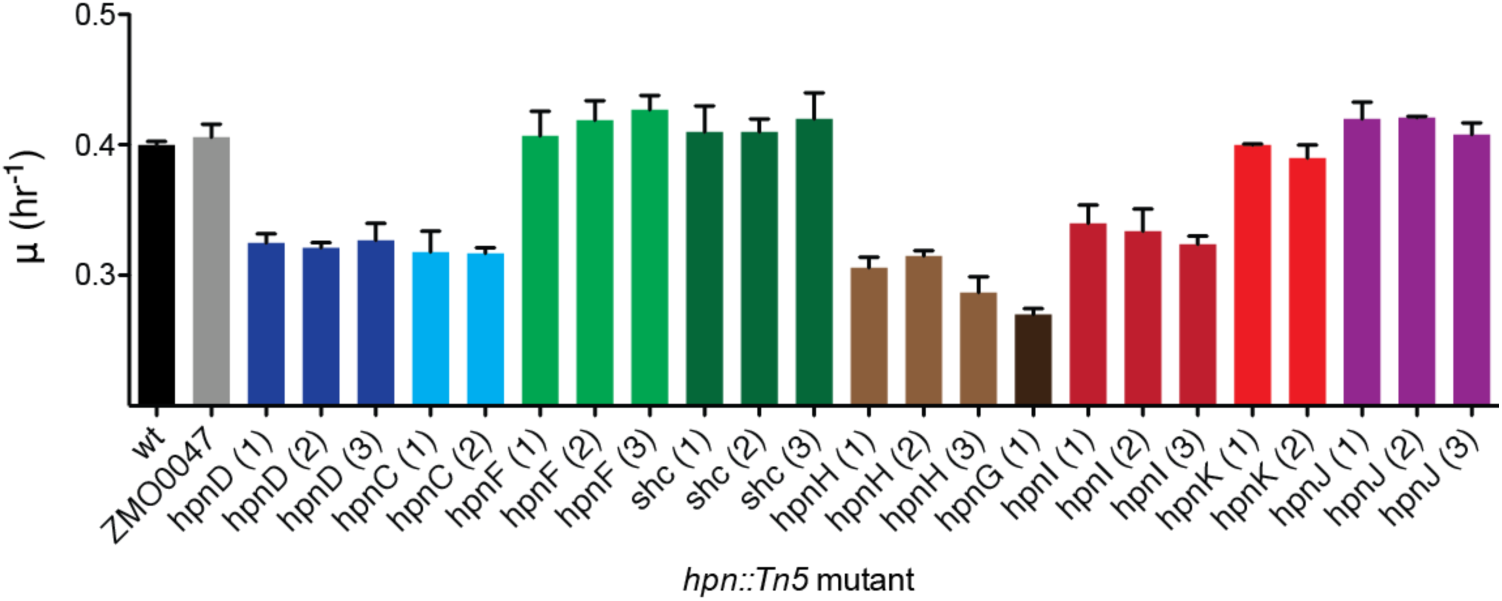
Redundant mutants of *hpn* genes, which feature Tn5 insertions in different sites, feature identical exponential growth rates (µ), indicating that all act similarly as loss of function mutations. Wild type cells (wt) and a kanamycin resistant control (ZMO0047, with a Tn5 insertion in a non-essential lysine exporter) are used as controls throughout this study. Because each redundant mutant featured similar fitness (e.g. *hpnD* (1) vs. *hpnD* (2)) only one of them was chosen for further genetic, chemical, and physiological analysis. Colors indicate mutants of the same gene. Error bars indicate SD.

**Fig. S2:**
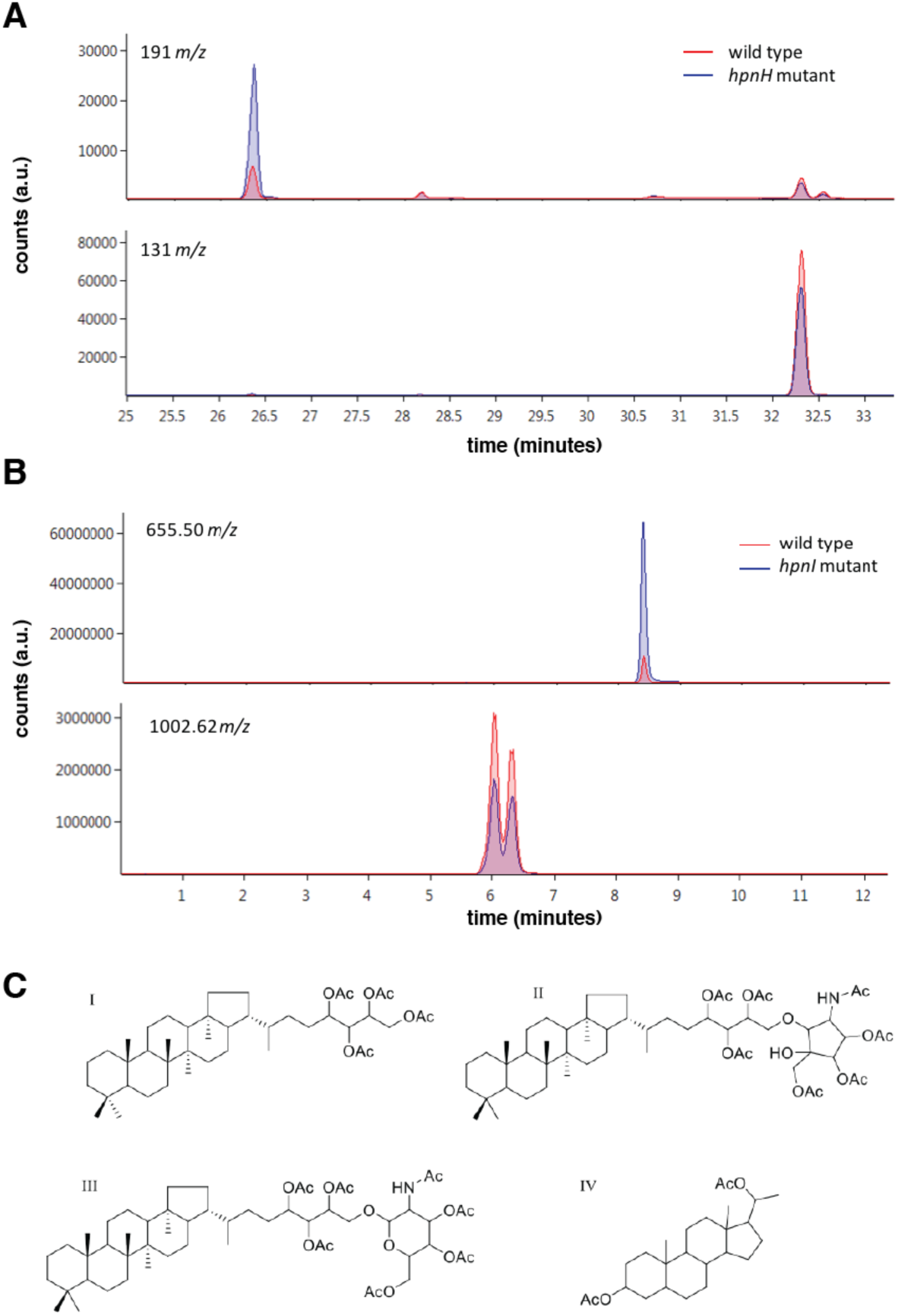
Sample chromatograms for hopanoid analyses methods used in this study. (**A**) GC-MS chromatograms showing the characteristic fragments for hopene (191 *m/z*, top) and diplopterol (131 *m/z*, bottom) for silylated lipids from a wild type *Z. mobilis* culture and one carrying a Tn5 insertion in *hpnH*. Because HpnH utilizes hopene as a substrate, the *hpnH* mutant features increased hopene, but not diploterol, levels. (**B**) APCI-LC-TOF-MS chromatograms showing the characteristic ions for acetylated lipids from a from a wild type *Z. mobilis* culture and one carrying a Tn5 insertion in *hpnI*. The mutant features reduced amounts of BHT (655.50 *m/z* representing [M+H-CH_3_COOH]^+^ for BHT, top) but reduced amounts of BHT glucosamine and BHT cyclitol ether (1002.62 *m/z* representing [M+H]^+^ for both species, bottom). The latter species are isomers, so they feature identical masses but can be separated by liquid chromatography. HpnI utilizes BHT as a substrate, so the accumulation of BHT is consistent with reduced copy numbers of *hpnI*. (**c**) Structure compounds analyzed by LC-APCI-TOF-MS; I-III represent acetylated hopanoid species, while IV is an acetylated internal standard.

**Fig. S3:**
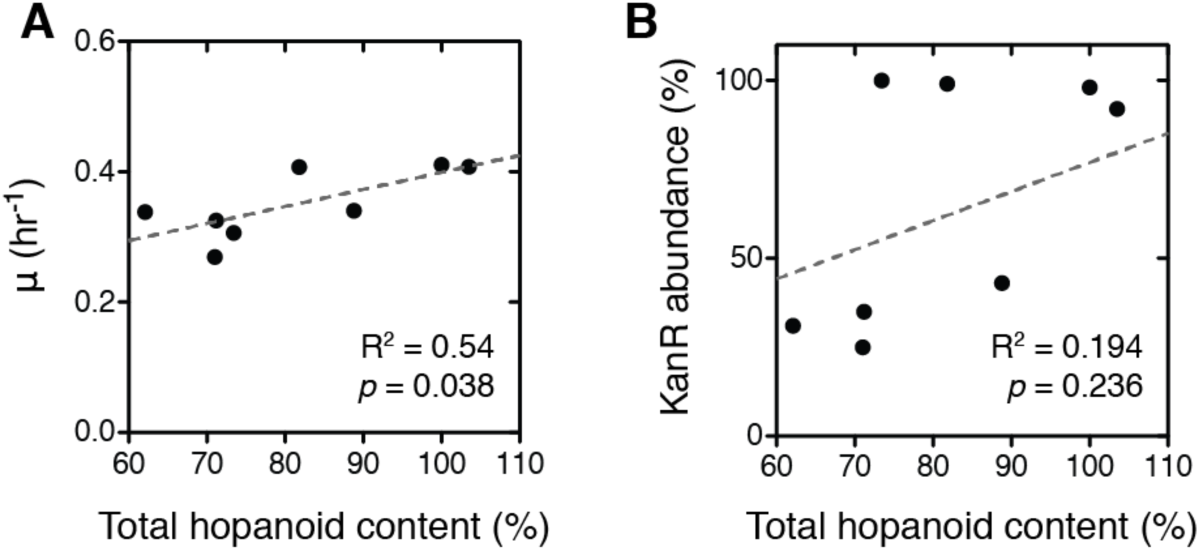
Growth rates (µ) of *hpn* mutants are significantly correlated with total hopanoid content (**A**) but not to the frequency of Tn5 alleles, as measured by sequencing read depth (**B**). Hopanoid content is presented as a % relative to wild type cells. *P*-values represent significance tests assessing if the slope of the linear regression is non-zero.

**Fig. S4:**
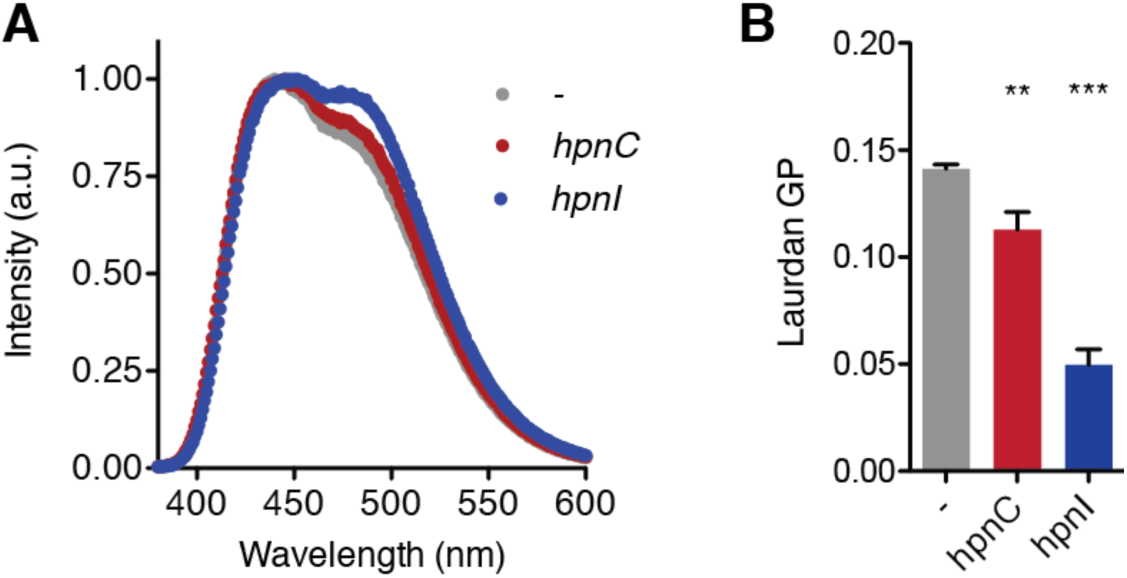
The emission spectra for Laurdan (excitation at 364 nm) incubated in liposomes assembled from extracted *Z. mobilis* lipids. Liposomes from control cells (-), featuring intact hopanoid compositions, exhibit a Laurdan emission spectrum that is blue shifted compared to those from cells with reduced (*hpnC*) or altered (*hpnI*) hopanoid levels. This emission difference, quantified by General Polarization (GP) values in the text, measures solvation of the membrane. Increased presence of polar solvent leads to relaxation of the excited dye, thereby red-shifting its emission in membranes that are less ordered near the head group-hydrocarbon interface. Error bars indicate mean +/-SD, n = 3 independent liposome preparations. Unpaired t-tests were used to assess significance against liposomes with intact hopanoids: **, p < 0.01; ***, p < 0.001.

